# Compartment-Specific Activation of the Proton-Sensor GPR65 is Uncoupled from Receptor Trafficking

**DOI:** 10.1101/2023.03.18.533272

**Authors:** Loyda M. Morales Rodríguez, Stephanie E. Crilly, Jacob B. Rowe, Daniel G. Isom, Manojkumar A. Puthenveedu

**Affiliations:** Department of Pharmacology, University of Michigan Medical School, Ann Arbor, United States; Cellular and Molecular Biology Program, University of Michigan Medical School, Ann Arbor, United States; The Department of Molecular and Cellular Pharmacology, University of Miami Miller School of Medicine, Miami, Florida, United States; Sylvester Comprehensive Cancer Center, University of Miami Miller School of Medicine, Miami, Florida, United States; Institute for Data Science and Computing, University of Miami Miller School of Medicine, Miami, Florida, United States

## Abstract

The canonical view of G protein-coupled receptor (GPCR) function is that receptor trafficking is tightly coupled to signaling. GPCRs remain on the plasma membrane (PM) at the cell surface until they are activated, after which they are desensitized and internalized into endosomal compartments. This canonical view presents an interesting context for proton-sensing GPCRs because they are more likely to be activated in acidic endosomal compartments than at the PM. Here we show that the trafficking of the prototypical proton-sensor GPR65 is fully uncoupled from signaling, unlike that of other known mammalian GPCRs. GPR65 internalized and localized to early and late endosomes, from where they signal at steady state, irrespective of extracellular pH. Acidic extracellular environments stimulated receptor signaling at the PM in a dose-dependent manner, although endosomal GPR65 was still required for a full signaling response. Receptor mutants that were incapable of activating cAMP trafficked normally, internalized, and localized to endosomal compartments. Our results show that GPR65 is constitutively active in endosomes, and suggest a model where changes in extracellular pH reprograms the spatial pattern of receptor signaling and biases the location of signaling to the cell surface.

## INTRODUCTION

For all known members of the physiologically and clinically important G protein-coupled receptor (GPCR) family (1,2), signaling and membrane trafficking are tightly coupled (3,4). GPCRs activated at the plasma membrane (PM) on the cell surface are rapidly desensitized and internalized into endosomal compartments, from where they can either recycle back to the PM or be degraded in the lysosome. This sorting determines the further responsiveness of cells to ligands (5,6). Endosomes also serve as signaling stations for many GPCRs. Importantly, the same signals originating from GPCRs in the plasma membrane versus endosomes can produce distinct downstream consequences (4,7–10). These observations have led to an emerging model that the GPCR signaling is spatially encoded, where the integrated GPCR response is a balance of both surface and internal signals, determined by rates of trafficking of receptors to and from the PM.

The relationship between trafficking and signaling is especially interesting for proton-sensing GPCRs, which are physiologically important but are understudied (11). These GPCRs are highly overexpressed in many cancers (12), and they have generated interest as potential therapeutic targets. The canonical view is that these receptors are activated by acidic extracellular environments, which are defining hallmarks of cancer and inflammation as well as many physiological processes (13–16). Unlike most canonical GPCRs which are desensitized to extracellular ligands upon endocytosis, endocytosis of proton-sensing GPCRs transports them to endosomal compartments that are acidic and more likely to activate these receptors. Considering the intrinsic signaling potential of proton sensing GPCRs in intracellular compartments, whether these receptors follow the tight coupling of trafficking and signaling that have been described for most known GPCRs, or whether they can be selectively activated in endosomal compartments, are unanswered questions that are fundamental to understanding how cells respond to pH.

Here we addressed these questions using the proton-sensing receptor GPR65 as a prototype. GPR65 is highly overexpressed in many solid tumors and are emerging as attractive targets to treat cancer, as they are thought to respond to acidic microenvironments and modify tumor signaling and the immune response. We show that GPR65 internalizes from the PM irrespective of extracellular pH, and that receptor internalization is required for a full cellular signaling response. Together, our findings show that GPR65 dynamically traffics to and signals from multiple cellular compartments, and that, unlike for most known GPCRs, activation of GPR65 is uncoupled from receptor trafficking. Characterizing proton-sensing receptor trafficking and signaling will improve our understanding of how these receptors integrate responses from multiple locations in the cell, and how these responses influence physiology and disease states.

## RESULTS

### GPR65 stimulates cAMP accumulation at neutral and acidic pH

We first determined the pH-dependent activation of GPR65, by measuring levels of the second messenger cAMP in HEK293 cells stably expressing GPR65. We used the cAMP biosensor GloSensor, which exhibits increased luminescence when bound to cAMP, to quantitatively measure cAMP levels (**Figure 1A**) (17,18). Upon exposure to proton concentrations from pH 6.0 to 8.0, GPR65-expressing cells display a dose-responsive elevation of cAMP levels (**Figure 1B and C**). This elevation in cAMP saturated at pH 6.4. Interestingly, cAMP levels were elevated at both acidic and neutral pH ranges. As negative controls, HEK293 cells not expressing GPR65 did not show a response (**(Figure 1D-E)**. Similarly, a GPR65 mutant where three key histidines, H10, H14 and H243, were mutated, which has been shown to not increase cAMP levels, did not show a cAMP response (**Figure 1D-E**) (13,19). This GPR65-mediated cAMP increase is similar to the prototypical Gs-coupled GPCR beta-2 adrenergic receptor (B2AR), which we used as a positive control for cAMP activation (**Figure 1D-E**). When cells expressing GPR65 were treated with a low concentration of protons (high pH), GloSensor luminescence rapidly decreased below that of vehicle/pH 7.0 (**Figure 1B and C, Supplemental figure 1B**). Both acute and chronic basic pH treatments decreased GPR65-mediated GloSensor luminescence (**Figure 1B, Supplemental figure 1B**). The absolute cAMP response and the changes were reduced substantially in HEK293 cells not expressing GPR65 (**Supplemental figure 1A-B**), indicating that the cAMP response we observed in GPR65-expressing cells were a result of GPR65 activation. In HEK293 cells, forskolin-induced cAMP responses did not change until pH 8.0, indicating that the dose response observed in GPR65-expressing cells was not a direct effect of pH on the sensor. Together, these data demonstrate that GPR65 increases cAMP levels at both acidic and neutral pH.

**Figure 1.**
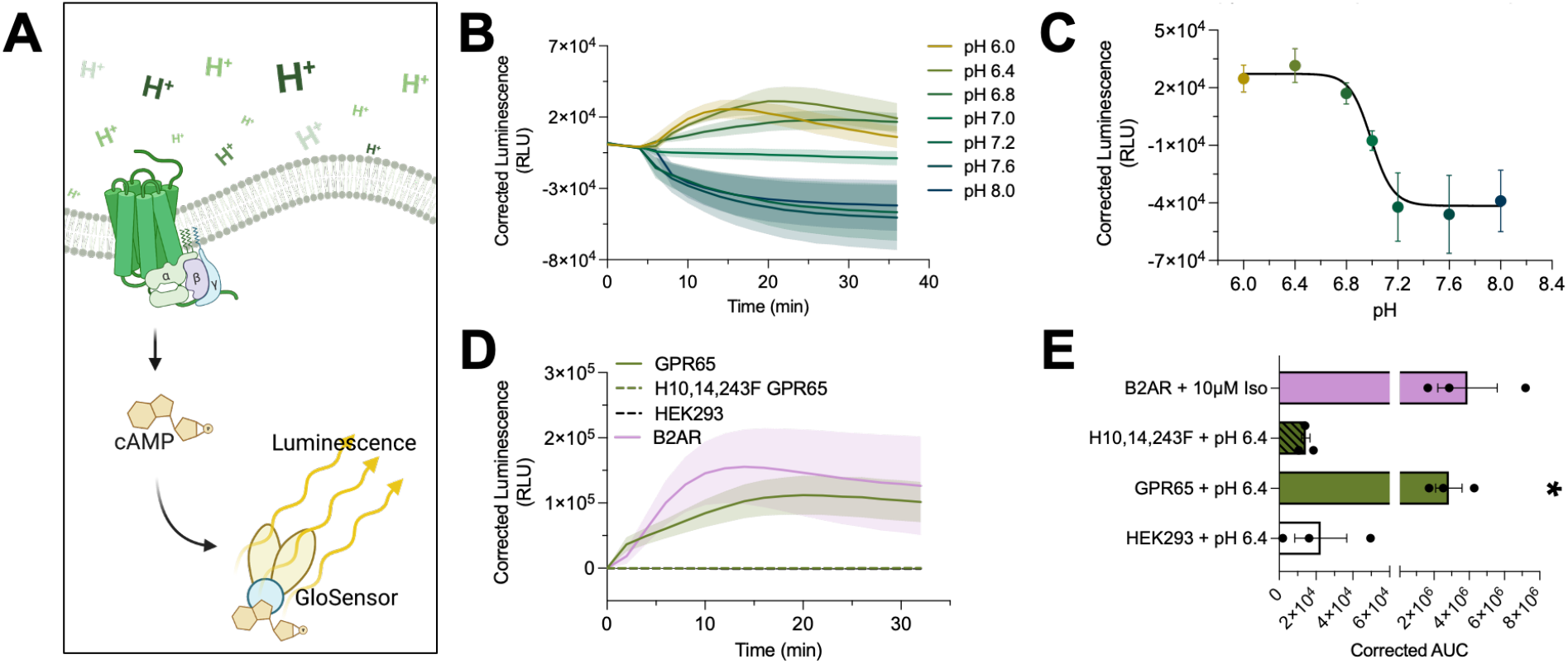
GPR65 increases cAMP levels at both neutral and acidic pH. (**A**) Schematic diagram of the GPR65-GloSensor assay. Proton-dependent activation of GPR65 stimulates production of intracellular cAMP. cAMP binding to the GloSensor luciferase produces luminescence directly correlated to increased cAMP levels. (**B**) GloSensor luminescence over time in GPR65-expressing cells treated with pH 6.0, 6.4, 6.8, 7.0, 7.2, 7.6 or 8.0 (n=3 biological replicates). GPR65 displays elevated cAMP levels at neutral and acidic pH. (**C**) Proton-dependent changes in intracellular cAMP levels were measured by a concentration-response curve (pH 6.0-8.0) in HEK293 cells stably expressing FLAG-GPR65, and GloSensor (n=3 biological replicates, Nonlinear regression fit [agonist] vs response-Variable slope). GPR65-mediated cAMP increase is saturable at pH 6.4. (**D**) GloSensor luminescence over time in HEK293 cells and HEK293 cells stably expressing FLAG-GPR65, FLAG-B2AR and FLAG-H10F, H14F, H243F-GPR65 mutant after addition of either pH 6.4 or Iso (n=3 biological replicates). Activation of GPR65 by pH 6.4 increases cAMP levels similar to positive control B2AR. (**E**) Bar graph of the area-under-curve (AUC) from figure 1D (n=3 biological replicates) (Ordinary one-way ANOVA, p<0.05). AUC of WT GPR65 is similar to positive control B2AR and significantly higher than negative controls.

### GPR65 internalizes from the plasma membrane irrespective of extracellular pH

Because GPR65 exhibited increased cAMP levels at a physiologically relevant pH range, we asked whether GPR65 was constitutively internalizing from the PM, and how the internalization compared to the prototypical B2AR. To test this, HEK293 cells expressing epitope-tagged FLAG-GPR65 were immunolabeled live (**Figure 2A**) and imaged by confocal microscopy after 10 minutes of labeling at 37°C. At neutral and basic pH, FLAG-GPR65 localizes to the PM and internal compartments (**Figure 2B and C**), suggesting that surface receptors internalized rapidly at these pH levels. When exposed to acidic pH, FLAG-GPR65 localization does not change noticeably and is again observed at the PM and internal compartments (**Figure 2B and C**). Quantification of the number of internal receptor spots shows a similar degree of internalization of FLAG-GPR65 exposed to pH 6.4-8.0 (**Figure 2C).** This agonist-independent internalization pattern is very different from the prototypical GPCR B2AR. At baseline, B2AR is localized primarily to the PM (**Figure 2B**). When exposed to a saturating concentration of agonist isoproterenol (10 μM), B2AR internalized and localized almost entirely to internal compartments (**Figure 2B and C**). Importantly, SNAP-tagged GPR65 displayed a similar internalization pattern to FLAG-tagged GPR65 (**Figure 2F and G**), indicating that the localization pattern was not a function of the FLAGtag. Together, these data demonstrate that GPR65 constitutively internalizes from the PM and localizes to internal compartments irrespective of pH.

**Figure 2.**
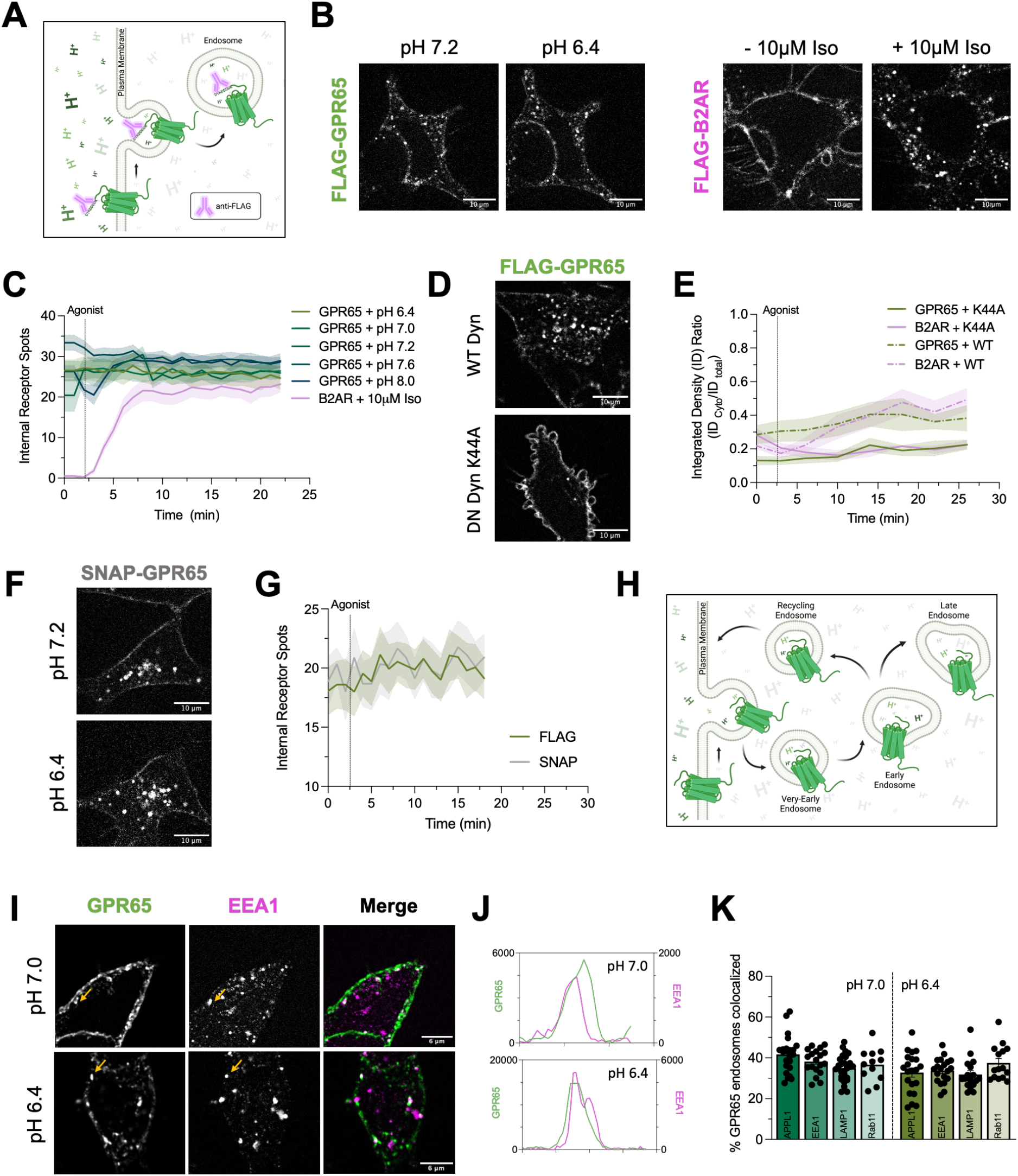
GPR65 internalization and endosomal localization is independent of extracellular pH. (**A**) Schematic diagram of the FLAG internalization assay. A fluorescently tagged antibody will bind the FLAG tag (peptide sequence DYKDDDDK) at the N-terminus of the receptor. Immunolabeling live cells with the tagged antibody will allow for visualization of receptors that started at the PM when cells were labeled. (**B**) FLAG-tagged GPR65-expressing cells exposed to pH 8.0 (left) and pH 6.4 (right) were immunolabeled live and imaged by confocal microscopy. Localization of FLAG-GPR65 does not change noticeably. B2AR-expressing cells were immunolabeled live and imaged by confocal microscopy. After addition of 10uM Iso, B2AR internalizes and localizes almost entirely to internal compartments. (**C**) Quantification of the number of internal receptor spots over time (FLAG-GPR65, n=15-25 cells/condition; FLAG-B2AR, n=16 cells) (symbol indicates mean, shading is SEM). FLAG-GPR65 localization remains constant even after pH 6.4 addition. (**D**) FLAG-GPR65 cells expressing either dominant-negative Dyn K44A or WT Dyn were immunolabeled live and imaged by confocal microscopy. FLAG-GPR65 cells expressing Dyn K44A exhibit decreased endocytosis while cells expressing WT Dyn displayed increased internalization and endocytosis. **(E)** Quantification of the ratio of internal fluorescence over total cell fluorescence over time (WT Dyn, n=20 cells each; Dyn K44A, n=20 cells each) (symbol indicates mean, shading is SEM). FLAG-GPR65 cells expressing Dyn K44A exhibit a lower internal fluorescence while cells expressing WT Dyn display increased internal fluorescence. (**F**) SNAP-tagged GPR65-expressing cells exposed to pH 7.2 (top) and pH 6.4 (bottom) were immunolabeled live and imaged by confocal microscopy. Localization of SNAP-GPR65 is similar to FLAG-GPR65 (scale bars= 10 μm). (**G**) Quantification of the number of internal receptor spots over time. The FLAG-GPR65 internalization pattern is conserved in SNAP-GPR65 (SNAP-GPR65, n=15 cells; FLAG-GPR65, n=15 cells) (symbol indicates mean, shading is SEM). (**H**) Schematic of the endocytic pathway that allows for trafficking and transfer of cargoes between membrane-bound compartments. Once a membrane protein is internalized, the protein is transferred from very-early (APPL1+) endosomes to early (EEA1+) endosomes. From EEA1 + endosomes, the protein could either be inserted back into the PM via recycling endosomes (like Rab11+ endosomes) or it could be targeted for degradation via late endosomes (like LAMP1 + endosomes). (**I**) Representative confocal images of HEK293 cells expressing FLAG-GPR65 exposed to pH 7.0 or pH 6.4 for 20 min prior to fixing cells and staining for EEA1 with a 488 secondary antibody (scale bar= 6 μm). Yellow arrows denote GPR65 endosomes that colocalize with EEA1. (**J**) Fluorescence linear profile plots of GPR65 and EEA1, measured by lines drawn across regions of the cell with GPR65 endosomes after treatment with pH 7.0 or pH 6.4 for 20 min. Plots show EEA1 immunofluorescence increases along with GPR65 in both pH 7.0 and pH 6.4. (**K**) Quantitation of the percentage of GPR65 containing endosomes that colocalize with each of the endosomal markers. For this, FLAG-GPR65 cells treated with either pH 7.0 or pH 6.4 for 20 min were fixed and processed for immunofluorescence with the noted markers. GPR65 localizes in APPL1, EEA1 Rab11 and Lamp1 positive endosomes irrespective of pH (n = 21-23, 19-20, 13-16, and 17-28 fields for APPL1, EEA1, Rab11, and Lamp1, respectively).

To directly confirm that the internal GPR65 was internalized from the PM, we inhibited endocytosis and tested whether this changed GPR65 localization. We expressed a dominant-negative (DN) dynamin (Dyn) mutant (K44A), which inhibits receptor-mediated endocytosis (20), in both FLAG-GPR65 and FLAG-B2AR cells, immunolabeled live, and imaged cells by confocal microscopy. Live imaging revealed FLAG-GPR65 cells expressing Dyn K44A exhibit decreased endocytosis while cells expressing WT Dyn displayed internalization and endocytosis (**Figure 2D**). Quantification of the ratio of internal fluorescence over total cell fluorescence throughout time revealed that GPR65 cells expressing Dyn K44A exhibited a lower internal fluorescence while cells expressing WT Dyn displayed increased internal fluorescence (**Figure 2D**). FLAG-B2AR cells expressing Dyn K44A, or WT Dyn and treated with isoproterenol (Iso) showed similar results to GPR65 cells expressing Dyn K44A and WT Dyn, respectively (**Figure 2E**). Together, these data demonstrate GPR65 internalizes from the PM in a dynamin-dependent manner irrespective of pH and localizes to endosomes.

We next identified the endosomal compartments to which GPR65 localized, and tested whether the localization pattern of GPR65 changed between neutral and acidic extracellular pH. We treated HEK293 cells expressing FLAG-GPR65 with pH 7.0 or pH 6.4 for 20 min, immunolabeled live, fixed, and stained cells with markers for the distinct compartments along the endosomal pathway: APPL1 (very early endosomes), EEA1 (early endosomes), Rab11 (recycling endosomes), and Lamp1 (lysosomes) (**Figure 2H**). Using the Imaris spot detection and colocalization analysis, we quantified the fraction of GPR65 spots that colocalized with each endosomal marker. At both pH 7.0 and pH 6.4, GPR65 localized to multiple compartments, specifically EEA1, APPL1, Rab11 and Lamp1 endosomes (**Figure 2I-K**). These results show that GPR65 is present in early and late endosomes across a wide pH range, which was surprising considering that GPCR activation and trafficking are usually highly integrated (4,7,21,22).

### Proton-dependent activation of GPR65 is uncoupled from receptor trafficking

Because GPR65 was localized in internal compartments across its whole signaling spectrum, we next directly asked whether internalization and endosomal localization of surface-labeled GPR65 required the receptor to be able to signal, by testing the trafficking of GPR65 mutants that were deficient in activating cAMP. In addition to the histidine mutant (H10F, H14F and H243F) described in **Figure 1**, which did not stimulate cAMP, we generated an independent GPR65 mutant deficient in cAMP signaling by mutating R112 in the DRY Motif, a conserved stretch of amino acids that governs GPCR activation and G protein coupling (23). To confirm that the GPR65 DRY Motif mutant R112A was deficient in cAMP signaling, the mutant was expressed in HEK293 cells expressing the GloSensor prior to pH 6.4 addition. At neutral pH, WT GPR65 exhibited increased luminescence while GPR65 R112A displays lower luminescence similar to HEK293 cells (**Figure 3A**). After exposure to pH 6.4, WT GPR65 displayed increased luminescence while GPR65 R112A and HEK293 cells did not increase cAMP levels in response to acidic pH (**Figure 3B**). Mutation of the arginine residue in the DRY Motif abolished the proton-dependent cAMP increase observed in WT GPR65 (**Figure 3A-C**). Strikingly, FLAG-R112A GPR65 and FLAG-H10, 14, 243F GPR65 mutants were both localized to intracellular endosomal structure, identical to WT FLAG-GPR65, when visualized via confocal microscopy (**Figure 3D-E**). Together, these results suggest that GPR65 activation was not required for GPCR internalization, and that GPR65 trafficking and signaling were uncoupled.

**Figure 3.**
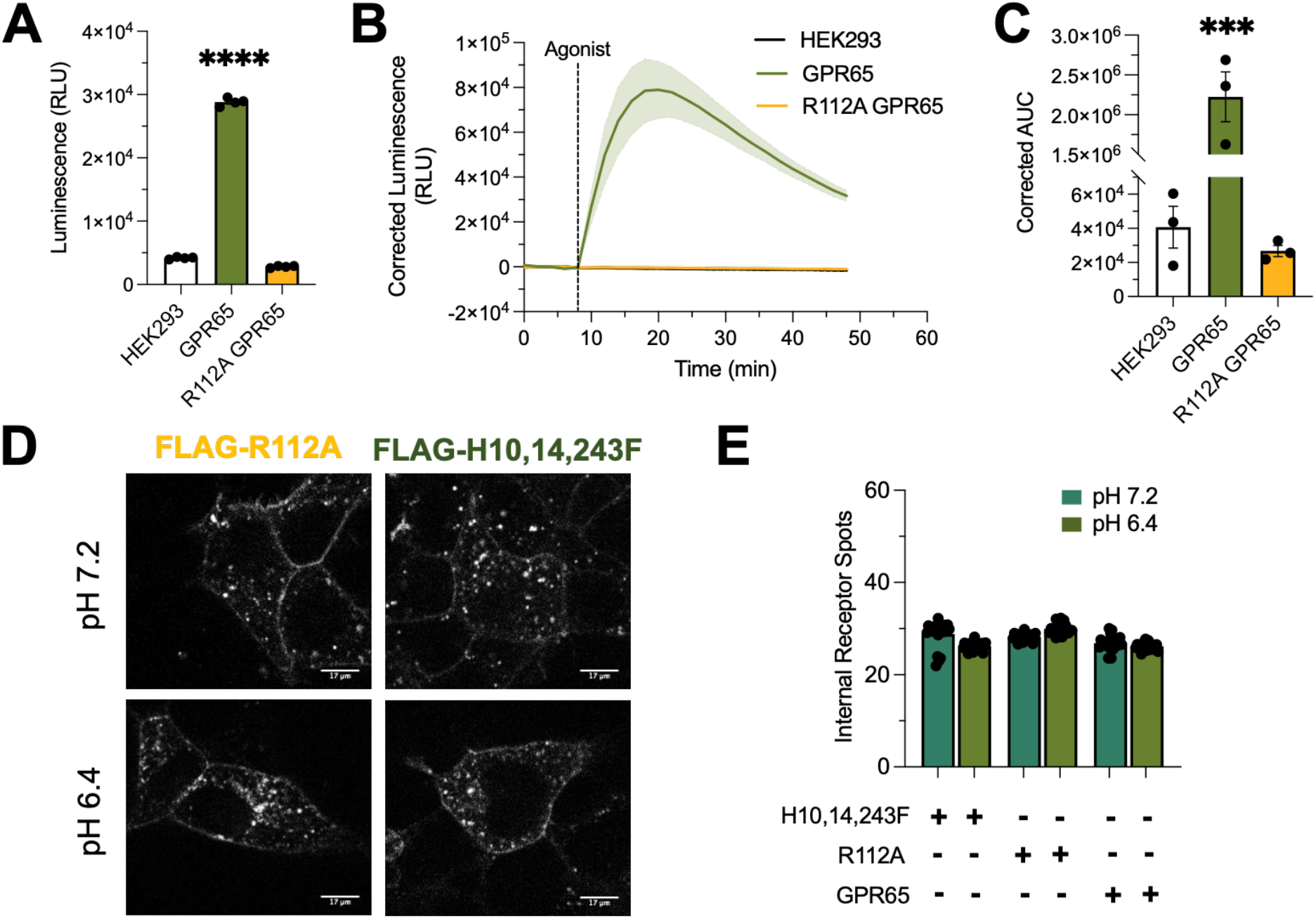
Activation of GPR65 is uncoupled from receptor trafficking. (**A**) Luminescence at pH 7.0 of GPR65-expressing cells compared to R112A-GPR65 DRY Motif mutant and HEK293 cells (n=4 biological replicates) (Ordinary one-way ANOVA, p<0.0001). At neutral pH, WT GPR65 exhibits increased luminescence while R112A-GPR65 displays lower luminescence similar to HEK293. (**B**) Corrected luminescence trace of GPR65-expressing cells compared to R112A-GPR65 DRY Motif mutant and HEK293 cells exposed to pH 6.4 (n=3 biological replicates). After exposure to pH 6.4, WT GPR65 cells display increased luminescence while R112A-GPR65 DRY Motif mutant and HEK293 cells did not increase cAMP levels in response to acidic pH. (**C**) Bar graph of the area-under-curve (AUC) from figure 3B (n=3 biological replicates) (Ordinary one-way ANOVA, p<0.05). AUC of WT GPR65 is significantly higher than R112A-GPR65 DRY Motif mutant and the negative control, HEK293 not expressing GPR65. (**D**) FLAG-R112A and FLAG-H10,14,243F GPR65-expressing cells exposed to pH 7.2 (left) and pH 6.4 (right) were immunolabeled live and imaged by confocal microscopy. Localization of FLAG-R112A GPR65 and FLAG-H10, 14, 243F GPR65 do not change noticeably upon pH 6.4 addition. (**E**) Quantification of the number of internal receptor spots over time (FLAG-H10, 14, 243F GPR65, n=25-32 cells; FLAG-R112A GPR65, n=22-29 cells; GPR65, n=15-25 cells) (symbol indicates mean, error bars are SEM). FLAG-R112A and H10,14,243F GPR65 localization is similar to WT GPR65.

### Internalized GPR65 contributes to whole-cell cAMP response

Localization of GPR65 to distinct intracellular compartments at steady-state raised the possibility that GPR65 was consistently active at acidic endosomes. To test this possibility, we pretreated cells with the endocytosis inhibitor dyngo-4a (**Figure 4A**) for 15 min, and measured cAMP signaling via GloSensor luminescence before and after pH 6.4 addition. Inhibition of GPR65 endocytosis reduced total cAMP response when compared to vehicle-treated cells (**Figure 4B and D**). This significant reduction in cAMP response of dyngo-4a treated GPR65-expressing cells is similar to B2AR cells exposed to dyngo-4a (**Figure 4B-D**), where part of the iso-activated cAMP response comes from endosomes (7,24). Together, these data suggest internalized GPR65 contributes to the persistent cAMP response observed in GPR65 expressing cells.

**Figure 4.**
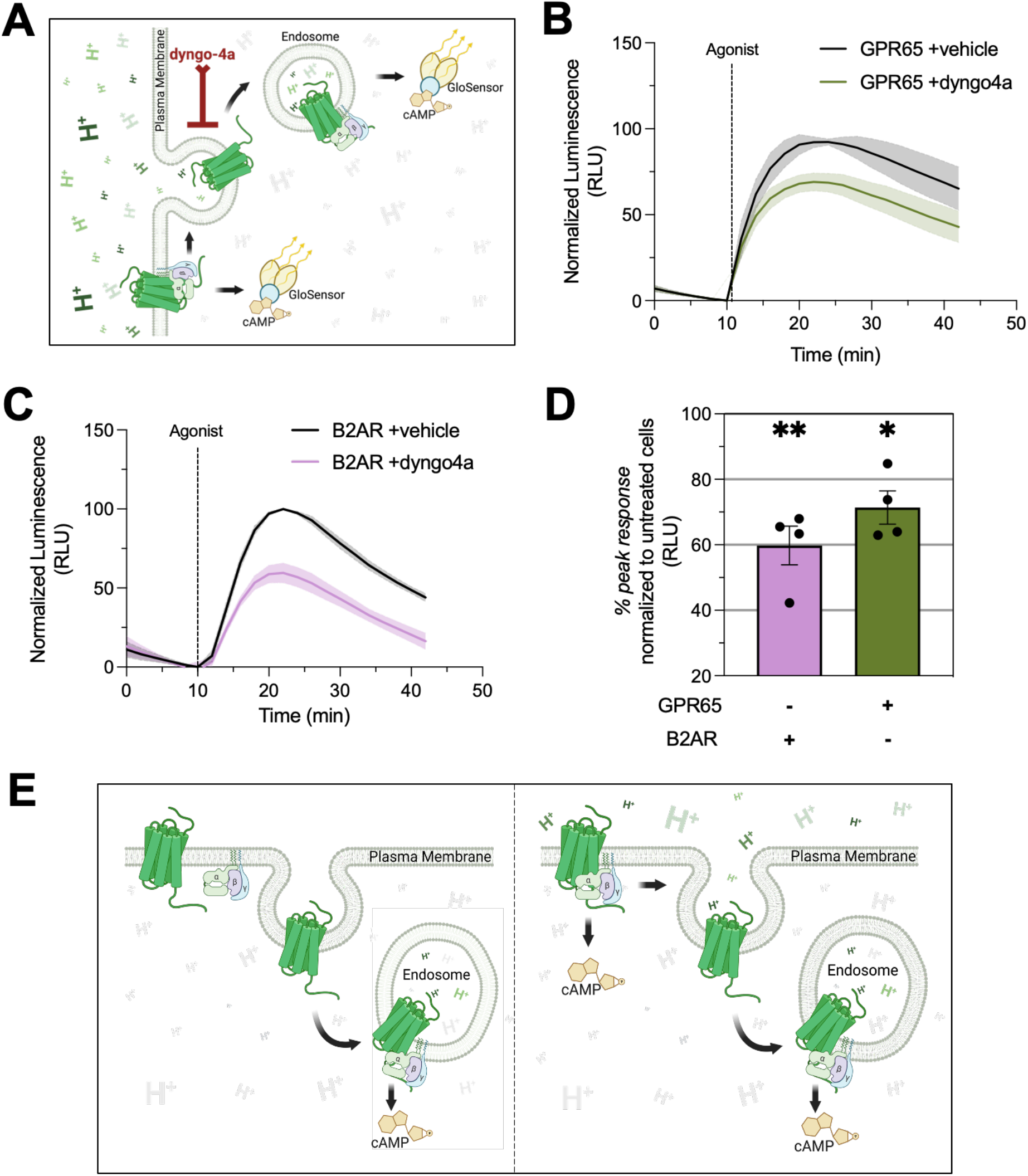
Internalized GPR65 contributes to cAMP response. (**A**) Proton-dependent activation of GPR65 stimulates production of intracellular cAMP from multiple cellular compartments. Treatment of GPR65-expressing cells with endocytosis inhibitor dyngo-4a will prevent internalization of the receptor leading to a decreased whole-cell cAMP output. (**B**) Corrected luminescence trace of GPR65-expressing cells pretreated with vehicle or dyngo-4a for 15 min before addition of pH6.4 (n=4 biological replicates). After exposure to pH6.4, WT GPR65 cells treated with dyngo-4a display lower luminescence and intracellular cAMP levels when compared to vehicle condition. (**C**) Corrected luminescence trace of B2AR-expressing cells pretreated with vehicle or dyngo-4a for 15 min before addition of 10uM Iso (n=4 biological replicates). Positive control B2AR cells treated with dyngo-4a display lower luminescence and intracellular cAMP levels when compared to vehicle condition. (**D**) Peak luminescence response of GPR65 and B2AR in cells pre-treated with dyngo-4a, normalized to cells without dyngo-4a (n=4 biological replicates) (Welch’s t test, p<0.05). WT GPR65 cells treated with dyngo-4a display a significantly lower whole-cell cAMP output than vehicle. This decrease in GPR65 cAMP output is similar to B2AR. **(E)** A model for location-switching of GPR65 signaling triggered by extracellular pH. WT GPR65 exhibits constitutive internalization and localization to internal compartments even at a lower proton concentration in the extracellular space, ie. in a ligand- and activation-independent manner. In the acidic environment of the endosome, endosomal GPR65 becomes activated and tonically increases cAMP levels. An acidic extracellular environment activates PM GPR65 further increasing cAMP levels.

## DISCUSSION

Here we show that GPR65 localizes to endosomal compartments and stimulates cAMP production from endosomes independently from extracellular pH changes. Surprisingly, GPR65 activity was not required for receptor endosomal localization. Our results show that endosomal GPR65 sets the basal cAMP tone, and with acidic activation of GPR65 at the plasma membrane further increasing cAMP levels. These two sources of cAMP likely result in distinct pools of cAMP with distinct cellular functions.

Our results suggest that, contrary to known examples of mammalian GPCRs, trafficking of GPR65 to endosomal compartments is fully uncoupled from receptor activation at the plasma membrane. Prototypical class A GPCRs, like B2AR, are activated by agonist at the PM inducing a cascade of events that cause receptors to internalize and localize to internal compartments. GPR65 localized in endosomes even at extracellular pH conditions that did not stimulate cAMP above baseline (**Figure 2**), indicating that GPR65 internalization is ligand- and activation-independent. Ligand-independent internalization has been reported for the viral GPCR US28, which is constitutively active and is localized to internal compartments (25–27). Similarly, cannabinoid receptors or delta opioid receptors, show relatively high basal activity in the absence of added external ligands, and therefore show higher internalization in the absence of activating ligands. In these cases, receptor internalization is tightly coupled to its activation state. When these receptors are inactivated either by inverse agonists or by mutations, receptor endocytosis is substantially inhibited (28,29). In contrast, mutations in the GPR65 DRY motif that completely block signaling have no effect on the endosomal distribution of GPR65 (**Figure 3 D, E**).

The uncoupling of GPR65 activation and endosomal localization suggests that GPR65 signaling is regulated differently from other GPCRs. Activation of most GPCRs by ligand binding, typically at the PM, switches receptors from an “off” state to an “on” state. The initial G protein activation on the PM induced by ligand binding is rapidly desensitized by phosphorylation and arrestin binding. After internalization, a second phase of G protein-mediated signaling is initiated on endosomes. Importantly, G protein signaling from the PM and endosomes, even though they activate the same second messengers such as cAMP, activate separate sets of genes downstream of signals (7,10,30). The physiological outcome of activating a receptor is the integrated response of these multiple phases of signaling, separated by time and space (31,32). Gi-coupled receptors, such as opioid receptors, can also be in active conformations on endosomes after ligand-dependent activation at the PM (21,33). For these known examples, because the ligand is extracellular, mechanisms exist to transport the ligand to the endosomes, either by transporters that allow movement of small ligands such as catecholamines across the membranes, or by trafficking mechanisms that internalize larger ligands such as peptides (21,32). Our results suggest that GPR65 activation differs from this canonical off-on response. GPR65 activation in endosomes is independent of specific ligand transport or trafficking mechanisms. Rather, the endosomal activation is primarily determined by constitutive sorting of the receptor to late endosomal compartments where low pH results in activation.

Based on this unique uncoupling of trafficking and signaling, we propose a model where GPR65 is constitutively active in endosomes, and where acidic extracellular environments, rather than globally turning receptors on, instead switch or bias the intensity, timing, and location of signaling to the PM (**Figure 4**). At a physiological pH of 7.4 GPR65 is inactive at the plasma membrane (**Figure 1C**). The steep dose-response in the pH range of 7.2 to 6.8 allows cells to rapidly switch signaling to the plasma membrane, which could induce rapidly variable signaling outcomes that depend on the conformational biases induced by the membrane environment at the plasma membrane vs. endosomes (34). Therefore, acidic environments such as those observed in solid tumors could convert “tonic” endosomal signaling by GPR65, which is physiologically beneficial for immune cells, where GPR65 is highly expressed, to “spikes” of surface cAMP signaling.

The model suggests interesting new aspects of how GPR65 activation regulates signaling in physiological systems such as in immune cells. A role for cAMP in modulating immune cells is well established, but the exact mechanisms by which cAMP regulates immune function, and whether endosomal and surface signals have different physiologically different outcomes, are still not fully understood. Signaling via cAMP has been studied mainly downstream of adrenergic receptors, which are expressed in both innate and adaptive immune cells (35,36). Activation of cAMP can drive the secretion of selected cytokines and reduce inflammatory responses and infiltration by innate immune cells. However, cAMP also inhibits immune cell activation and proliferation, chemokine-dependent cell migration, and secretion of other cytokines and interferons (36,37), which could collectively inhibit the effectiveness of immune cells in tumor clearance. How sensitive these multiple effects are to precise levels of cAMP is not fully known. Immune cells, as they infiltrate different environments are exposed to different extracellular pH, like in the acidic environment in solid tumors. It is possible that the baseline level of tonic cAMP signaling, via constitutive GPR65 signaling from endosomes, is critical for maintenance of normal function of immune cells, and that spikes of surface signaling via adrenergic agonists and GPR65 could cause rapid changes in immune cell function, depending on the precise cell type and immune environment.

Overall, our findings provide a new perspective on GPR65 function that is critical to understand physiological proton-sensing. GPR65 and the family of proton-sensing receptors have been less studied compared to most other families of GPCRs (38). Although it is clear that GPCRs signal from intracellular compartments and that signaling from each compartment could have specific physiological roles, these specific roles, including for GPR65, still have to be fully elucidated. In this context, our study underlines the importance of understanding how the trafficking and signaling of these highly relevant receptors are regulated, and how the uncoupling of these two aspects is important for the role of these receptors in physiology and disease.

## MATERIALS AND METHODS

### Cell culture and transfection

Cell lines used were validated, and cells were purchased from ATCC. Cells in the lab were routinely tested for mycoplasma contamination, and only uncontaminated cells were used and maintained at 37°C with 5% CO_2_. Stable clonal HEK293 cells expressing either GPR65, H10,14,243F GPR65 or B2AR N-terminally tagged with FLAG were cultured in DMEM high glucose (Cytiva, SH3024301) supplemented with 10% fetal bovine serum (FBS; Gibco, 26140079). Stable cell lines expressing one of the constructs were generated using Geneticin (Gibco, #10131035) as selection reagent. All stable cell line plasmid transfections were conducted with Effectene (Qiagen, #301425) as per manufacturer’s instructions. HEK293 cells were also transiently transfected with GPR65 or R112A GPR65 fused to Flag on its N-terminus using Effectene as per manufacturer’s protocol.

### DNA constructs

FLAG-GPR65 construct consists of an N-terminal signal sequence followed by a FLAG tag followed by the human GPR65 sequence in a pcDNA3.1 vector backbone. To create SNAP-GPR65, the receptor sequence was amplified from the FLAG-GPR65 construct by PCR with compatible cut sites (BamHI and XbaI) and ligated into a pcDNA3.1 vector containing an N-terminal sequence, followed by a SNAP tag. FLAG-H10,14,243F GPR65 and FLAG-R112A GPR65 DRY mutant were created with a full-length receptor sequence gene block from Integrated DNA Technologies (IDT) with restriction sites (AgeI and XbaI) compatible to FLAG-GPR65 vector backbone and ligated into a pcDNA3.1 vector containing an N-terminal sequence, followed by a FLAG tag. FLAG-B2AR construct was described previously (Bowman et al., 2016). WT Dyn and Dyn K44A were gifts from Adam Linstedt. pcDNA3.1 empty vector was a gift from Drs. Alan Smrcka and Hoa Phan.

### Reagents

Leibovitz L15 imaging medium (Gibco, #21083-027) was used as the vehicle (pH 7.0) to deliver the desired pH since the buffered medium covers a wide pH range. The pH was adjusted by adding either 0.1 M HCl (Fisher Scientific, A144S-500) or 1 M NaOH (Fisher Scientific, #S318-500) and measured using pH test strips (Fisher Scientific, #13-640-502). Isoproterenol (Iso, #I5627) was purchased from Sigma Aldrich and used at 10μM from a 10 mM frozen stock. Dyngo-4a was purchased from ApexBio (#B5997), dissolved in DMSO (Fisher Scientific, #BP231-100) and used at 40μM. Mouse anti-FLAG M1 monoclonal antibody (Sigma Aldrich, #F3040) conjugated to Alexa 647 (Invitrogen, #A20173) and SNAP-surface dye 647 (NEB, #S9102S) were purchased from Sigma Aldrich, Invitrogen and New England BioLabs, respectively. anti-APPL1 (1:200; CST, D83H4, #3858S), anti-EEA1 (1:50; CST, C45B10, #3288), anti-Rab11 (1:50; CST, C45B10, #3288) or anti-LAMP1 (1:100; CST, D2D11 XP, #9091) rabbit monoclonal antibodies were purchased from Cell Signaling Technology (CST). Alexa 488 goat anti-rabbit secondary antibody (1:1000; #A11008) was bought from Invitrogen.

### GloSensor cAMP assay

HEK293 cells (5-7 × 10^4^ cells per well) were plated in a 96-well plate (Costar Corning, #3917) coated with poly-D-lysine (Sigma Aldrich, #P6407) to allow for adherence of cells. The following amounts of DNA were used per well: 60 ng of pGloSensor-22F cAMP plasmid (Promega, E2301), 100 ng of receptor or empty vector (control, pCDNA3.1^+^). Reverse transfection was performed using Effectene. Twenty-four hours after transfection, cells were washed once with Leibovitz L15 medium (Gibco, #21083-027), and 100 μl of 500μ g/mL D-luciferin (Goldbio, LUCK-1G) in Leibovitz’s L-15 medium was added for 2 hours at room temperature. Luminescence was measured for 30-50 min using a Varioskan LUX multimode microplate reader. For basic pH experiments, cells were treated with 5 μM Forskolin (Sigma Aldrich, #F3917). Raw luminescence values or the values corrected to baseline before acute changes in pH are noted as described.

### Live cell imaging

HEK293 cells were plated onto 25mm coverslips (Electron Microscopy Sciences, #50949050) coated with poly-D-lysine (Sigma Aldrich, #P6407) to allow for adherence. Two days later, cells were labeled with M1-647 antibody (1:1000) for 10 min and imaged in Leibovitz L15 imaging medium at 37°C in a CO_2_-controlled imaging chamber, using a spinning disk confocal microscope (Andor, Belfast, UK) and a 60× objective. Confocal images were acquired with an iXon +897 electron-multiplying charge-coupled device camera (Andor, Belfast, UK) and solid-state lasers of 488 nm or 647 nm.

### Immunofluorescence of endosomal markers

HEK293 cells stably expressing FLAG-GPR65 were plated to poly-d-lysine (Sigma Aldrich) coated 12 mm glass coverslips (Fisher Scientific, #1254580P) and grown for 24-48 hr at 37 °C. Cells were labeled with M1 647 antibody (1:1000) for 10 min, exposed to either pH 7.0 or pH6.4 for 20min at 37°C, then fixed with 4% formaldehyde (FB002, Invitrogen) for 20 min at room temperature. Cells were rinsed with wash solution (PBS containing 1.25mM calcium chloride, 1.25mM magnesium chloride, with 5% FBS, 5% 1M glycine) twice and then blocked in PBS containing 1.25mM calcium chloride, 1.25mM magnesium chloride, with 5% FBS, 5% 1M glycine, and 0.75% Triton X-100. After, FLAG-GPR5 cells were incubated with either rabbit anti-APPL1 (1:200; CST, D83H4, #3858S), anti-EEA1 (1:50; CST, C45B10, #3288), anti-Rab11 (1:50; CST, C45B10, #3288) or anti-LAMP1 (1:100; CST, D2D11 XP, #9091) endosomal marker antibodies for 1hr. Cells were washed three times with PBS containing calcium and magnesium and then labeled with Alexa 488 goat anti-rabbit secondary antibody (1:1000; Invitrogen, #A11008) in a blocking buffer for 1 hr. Cells were washed three times for 5 min and coverslips were mounted onto glass slides (Fisher Scientific, #12550123) using Prolong Diamond Antifade Mountant (Invitrogen, #P36961). Confocal imaging of cells was performed using a spinning disk confocal microscope (Andor) and 100× objective. Representative images were taken across 10–20 fields per condition.

### Image analysis and quantification

Stacks and time-lapse images were collected as TIFF images and analyzed with either FIJI or Imaris (Schindelin et al., 2012). We quantified intracellular receptor in two ways. We determined the total number of receptor spots in the cytoplasm of cells, using the Imaris software (Andor) spot’s function. We also analyzed images with FIJI and quantified receptor fluorescence in a region of interest corresponding to the cytosplasm of the cell as a fraction of total receptor fluorescence. For the endosomal colocalization analysis, we acquired the percent colocalization of receptor spots with the endosomal marker over the total number of receptor-positive endosomes via the Imaris software (Andor) spot’s function and the MATLAB-based colocalize spots extension. Statistical tests and graphs were generated using Prism 9 (GraphPad Software).

## ACKNOWLEDGEMENTS

We thank Dr. Hoa Phan, and Dr. Alan Smrcka for valuable feedback on the project and writing, key reagents and equipment. We thank Dr. Carole Parent, Dr. Maria Castro, and Dr. Wenjing Wang for their valuable feedback on this project. We also thank Yating (Christina) Zheng and Dr. Adam Courtney for generously providing key reagents and equipment. LMR was supported by the National Science Foundation Graduate Research Fellowship under Grant DGE 1256260. MAP was supported by NIH GM117425, DA055026, and National Science Foundation (NSF) grant 1935926.

## SUPPLEMENTAL INFORMATION

### Supplemental figures

**Supplemental Figure 1.**
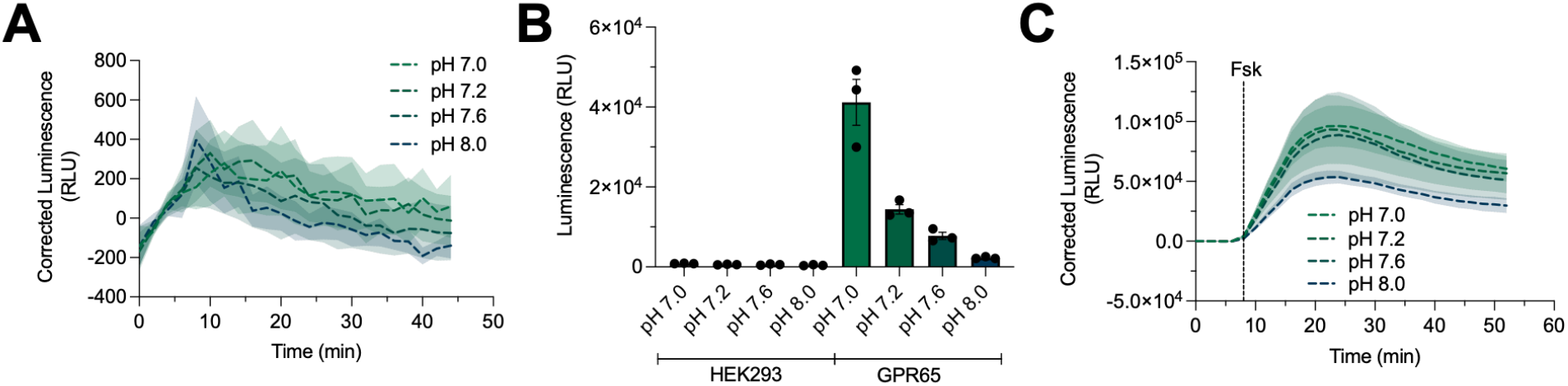
pH-sensitive cAMP activation in HEK cells is specific to GPR65 expression. (**A**) GloSensor luminescence trace in plain HEK293 cells exposed to a pH range from pH 7.0, to 8.0, after baseline readings (n=3 biological replicates). Exposure to a pH range from pH 7.0 to 8.0 over time does not significantly affect luminescence readings in HEK293 cells not transfected with GPR65 (untransfected). (**B**) GloSensor luminescence after two hours of neutral and basic pH (pH7.2, 7.6 and 8.0) pretreatment in HEK293 and GPR65-expressing HEK293 cells (n=6 biological replicates). Luminescence in GPR65-expressing cells depends on proton availability. (**C**) Forskolin (Fsk)-stimulated cAMP accumulation in untransfected HEK293 chronically treated with a pH range from 7.0 to 8.0 (n=3 biological replicates). Although a slight decrease is observed at pH 8.0, Fsk-induced GloSensor luminescence in untransfected HEK293 does not change in the dose-responsive pH range up to 7.6, showing that the sensor was not intrinsically sensitive to pH changes.

## REFERENCES

1. Pierce KL, Premont RT, Lefkowitz RJ. Seven-transmembrane receptors. Nat Rev Mol Cell Biol. 2002;3(9):639–50.

2. Sriram K, Insel PA. G protein-coupled receptors as targets for approved drugs: How many targets and how many drugs? Mol Pharmacol. 2018;93(4):251–8.

3. Sorkin A, Von Zastrow M. Endocytosis and signalling: Intertwining molecular networks. Nat Rev Mol Cell Biol. 2009;10(9):609–22.

4. Vilardaga J-P, Jean-Alphonse FG, Gardella TJ. Endosomal generation of cAMP in GPCR signaling. Nat Chem Biol. 2014;32(1):41–52.

5. Lobingier BT, von Zastrow M. When trafficking and signaling mix: How subcellular location shapes G protein-coupled receptor activation of heterotrimeric G proteins. Traffic. 2019;20(2):130–6.

6. Weinberg ZY, Puthenveedu MA. Regulation of G protein-coupled receptor signaling by plasma membrane organization and endocytosis. Traffic. 2019;20(2):121–9.

7. Bowman SL, Shiwarski DJ, Puthenveedu MA. Distinct G protein-coupled receptor recycling pathways allow spatial control of downstream G protein signaling. J Cell Biol. 2016;214(7):797–806.

8. Irannejad R, Tomshine JC, Tomshine JR, Chevalier M, Mahoney JP, Steyaert J, et al. Conformational biosensors reveal adrenoceptor signalling from endosomes. 2013;495(7442).

9. Sutkeviciute I, Vilardaga JP. Structural insights into emergent signaling modes of G protein-coupled receptors. J Biol Chem. 2020;295(33):11626–42.

10. Tsvetanova NG, Irannejad R, Von Zastrow M. G protein-coupled receptor (GPCR) signaling via heterotrimeric G proteins from endosomes. J Biol Chem. 2015;290(11):6689–96.

11. Silva PHI, Camara NO, Wagner CA. Role of proton-activated G protein-coupled receptors in pathophysiology. Am J Physiol - Cell Physiol. 2022;323(2):C400–14.

12. Insel PA, Sriram K, Salmerón C, Wiley SZ. Proton-sensing G protein-coupled receptors: Detectors of tumor acidosis and candidate drug targets. Future Med Chem. 2020;12(6):523–32.

13. Ludwig M-G, Vanek M, Guerini D, Gasser J, Jones CE, Junker U, et al. Proton-sensing G-protein-coupled receptors. Nature. 2003;425(6953):93–8.

14. Erra Díaz F, Dantas E, Geffner J. Unravelling the Interplay between Extracellular Acidosis and Immune Cells. 2018 [cited 2022 Apr 18]; Available from: https://doi.org/10.1155/2018/1218297

15. Kato Y, Ozawa S, Miyamoto C, Maehata Y, Suzuki A, Maeda T, et al. Acidic extracellular microenvironment and cancer. Cancer Cell Int [Internet]. 2013;13(1):1. Available from: Cancer Cell International

16. Okajima F. Regulation of inflammation by extracellular acidification and proton-sensing GPCRs. Cellular Signalling. 2013.

17. Fan F, Binkowski BF, Butler BL, Stecha PF, Lewis MK, Wood K V. Novel genetically encoded biosensors using firefly luciferase. ACS Chem Biol. 2008;3(6):346–51.

18. Wang FI, Ding G, Ng GS, Dixon SJ, Chidiac P. Luciferase-based GloSensor™ cAMP assay: Temperature optimization and application to cell-based kinetic studies. Methods [Internet]. 2021;(October 2021). Available from: https://doi.org/10.1016/j.ymeth.2021.10.009

19. Wang JQ, Kon J, Mogi C, Tobo M, Damirin A, Sato K, et al. TDAG8 is a proton-sensing and psychosine-sensitive G-protein-coupled receptor. J Biol Chem. 2004;279(44):45626–33.

20. Altschuler Y, Barbas SM, Terlecky LJ, Tang K, Hardy S, Mostov KE, et al. Redundant and distinct functions for dynamin-1 and dynamin-2 isoforms. J Cell Biol. 1998;143(7):1871–81.

21. Stoeber M, Jullie D, Laeremans T, Steyaert J, Schiller PW, Manglik A, et al. A genetically encoded biosensor reveals location bias of opioid drug action. 2018;1–27.

22. Thomsen ARB, Jensen DD, Hicks GA, Bunnett NW. Therapeutic Targeting of Endosomal G-Protein-Coupled Receptors. Trends Pharmacol Sci. 2018;176(5):139–48.

23. Rovati GE, Capra V, Neubig RR. The highly conserved DRY motif of class A G protein-coupled receptors: Beyond the ground state. Mol Pharmacol. 2007;71(4):959–64.

24. Tsvetanova NG, von Zastrow M. Spatial encoding of cyclic AMP signaling specificity by GPCR endocytosis. Nat Chem Biol. 2014;10(12):1061–5.

25. Casarosa P, Bakker RA, Verzijl D, Navis M, Timmerman H, Leurs R, et al. Constitutive signaling of the human cytomegalovirus-encoded chemokine receptor US28. J Biol Chem [Internet]. 2001;276(2):1133–7. Available from: http://dx.doi.org/10.1074/jbc.M008965200

26. Fraile-Ramos A, Kledal TN, Pelchen-Matthews A, Bowers K, Schwartz TW, Marsh M. The human cytomegalovirus US28 protein is located in endocytic vesicles and undergoes constitutive endocytosis and recycling. Mol Biol Cell. 2001;12(6):1737–49.

27. Fraile-Ramos A, Pelchen-Matthews A, Kledal TN, Browne H, Schwartz TW, Marsh M. Localization of HCMV UL33 and US27 in endocytic compartments and viral membranes. Traffic. 2002;3(3):218–32.

28. Leterrier C, Lainé J, Darmon M, Boudin H, Rossier J, Lenkei Z. Constitutive activation drives compartment-selective endocytosis and axonal targeting of type 1 cannabinoid receptors. J Neurosci. 2006;26(12):3141–53.

29. Gendron L, Cahill CM, Zastrow M Von, Schiller PW, Pineyro G. Molecular Pharmacology of d-Opioid Receptors. 2016;324876(July):631–700.

30. Tsvetanova NG, Trester-zedlitz M, Newton BW, Peng GE, Jeffrey R, Jimenez-morales D, et al. Endosomal cAMP production broadly impacts the cellular phosphoproteome. 2021;(Iii):1–19.

31. Crilly SE, Ko W, Weinberg ZY, Puthenveedu MA. Conformational specificity of opioid receptors is determined by subcellular location irrespective of agonist. Elife. 2021;10:1–20.

32. Irannejad R, Pessino V, Mika D, Huang B, Wedegaertner PB, Conti M, et al. Functional selectivity of GPCR-directed drug action through location bias. Nat Chem Biol. 2017;13(7):799–806.

33. Kunselman JM, Gupta A, Gomes I, Devi LA, Puthenveedu MA. Compartment-specific opioid receptor signaling is selectively modulated by different dynorphin peptides. Elife. 2021;10:1–17.

34. Wingler LM, Lefkowitz RJ. Conformational Basis of G Protein-Coupled Receptor Signaling Versatility. Trends Cell Biol [Internet]. 2020;30(9):736–47. Available from: https://doi.org/10.1016/j.tcb.2020.06.002

35. Guereschi MG, Araujo LP, Maricato JT, Takenaka MC, Nascimento VM, Vivanco BC, et al. Beta2-adrenergic receptor signaling in CD4+ Foxp3+ regulatory T cells enhances their suppressive function in a PKA-dependent manner. Eur J Immunol. 2013;43(4):1001–12.

36. Padro C. J, Sanders V. M. Neuroendocrine regulation of inflammation. Semin Immunol. 2014;26(5):357–68.

37. Huang S, Apasov S, Koshiba M, Sitkovsky M. Role of A2a extracellular adenosine receptor-mediated signaling in adenosine-mediated inhibition of T-cell activation and expansion. Blood [Internet]. 1997;90(4):1600–10. Available from: http://dx.doi.org/10.1182/blood.V90.4.1600

38. Roth BL, Kroeze WK. Integrated approaches for genome-wide interrogation of the druggable non-olfactory G protein-coupled receptor superfamily. J Biol Chem. 2015;290(32):19471–7.

