## Supplemental Figure 1 for "Compartment-Specific Activation of the Proton-Sensor GPR65 is Uncoupled from Receptor Trafficking"

**SUPPLEMENTAL INFORMATION**

**Supplemental figures:**

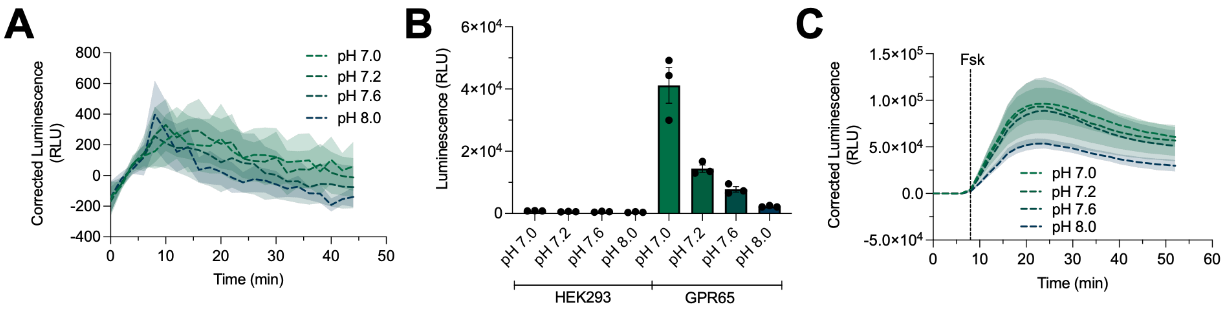


**Supplemental Figure 1. pH-sensitive cAMP activation in HEK cells is specific to GPR65 expression.** (**A**) GloSensor luminescence trace in plain HEK293 cells exposed to a pH range from pH 7.0, to 8.0, after baseline readings (n=3 biological replicates). Exposure to a pH range from pH 7.0 to 8.0 over time does not significantly affect luminescence readings in HEK293 cells not transfected with GPR65 (untransfected). (**B**) GloSensor luminescence after two hours of neutral and basic pH (pH7.2, 7.6 and 8.0) pretreatment in HEK293 and GPR65-expressing HEK293 cells (n=6 biological replicates). Luminescence in GPR65-expressing cells depends on proton availability. (**C**) Forskolin (Fsk)-stimulated cAMP accumulation in untransfected HEK293 chronically treated with a pH range from 7.0 to 8.0 (n=3 biological replicates). Although a slight decrease is observed at pH 8.0, Fsk-induced GloSensor luminescence in untransfected HEK293 does not change in the dose-responsive pH range up to 7.6, showing that the sensor was not intrinsically sensitive to pH changes.
